# Pro-autoimmune allele of tyrosine phosphatase, PTPN22, enhances tumor immunity

**DOI:** 10.1101/2021.03.17.435898

**Authors:** Robin C. Orozco, Kristi Marquardt, Kerri Mowen, Linda A. Sherman

## Abstract

The 1858C>T allele of the tyrosine phosphatase *PTPN22 (*causing amino acid substitution R620W in encoded protein Lyp) is present in 5-10% of the North American population and is strongly associated with numerous autoimmune diseases. Although much research has been done to define how this allele potentiates autoimmunity, the influence *PTPN22* and its pro-autoimmune allele has in tumor immunity is poorly defined. To interrogate the role this allele may have in the anti-tumor immune response, we used CRISPR/Cas9 to generate mice in which the ortholog of Lyp, PEP, is mutated at position 619 to produce the relevant pro-autoimmune mutation (R619W). Results of this study show that mice homozygous for this alteration (PEP-619WW) resist tumor growth as compared with wildtype mice. Consistent with these results, tumors from PEP-619WW mice have more CD45 infiltrates containing more activated CD8 T cell and CD4 T cells. Additionally, there are more cDC1 cells and less MDSCs in tumors from PEP-619WW mice. Interestingly, the tumor infiltrating PEP-619WW cDC1s have decreased PD-L1 expression compared to cDC1s from PEP-WT mice. Taken together, our data show that the pro-autoimmune allele of *Ptpn22* drives a strong anti-tumor response in innate and adaptive immune cells resulting in superior control of tumors.

## Introduction

Allelic variation in genes associated with regulating immune responses potentially impact an individual’s response to foreign and self-antigens. A minor allele of the protein tyrosine phosphatase non-receptor-type 22 (*PTPN22*) has been extensively studied for its contribution to autoimmunity (1-3). Genome-wide association studies identified a tryptophan-encoding common allelic variant, at amino acid 620 (R620W), that is associated with an increased risk of multiple autoimmune diseases such as Type 1 Diabetes (T1D), rheumatoid arthritis (RA), and systemic lupus erythematosus (SLE) (3-12). This allele is present in 5-15% of the North American and European populations and is considered the highest non-HLA autoimmune risk allele variant (2, 3). Therefore, understanding how *PTPN22* and its pro-autoimmune allele, regulates variation in immune cell functions is of great importance.

In humans, *PTPN22* encodes the protein Lymphoid tyrosine phosphatase (Lyp), the expression of which is confined to bone marrow derived cells and thus expressed in all immune cells (4). To better understand the role of this gene, researchers have often employed *Ptpn22* knock-out mice, which are deficient in expression of the Lyp ortholog, PEST-domain enriched protein or PEP (PEP-null) (13). These studies have shed light on the role of Lyp/PEP in multiple immune cell types. In lymphocytes, Lyp/PEP tempers T cell receptor (TCR) and B cell receptor (BCR) signaling through dephosphorylation of Src kinases (13, 14). Binding partners enabling such activity include TRAF3 and CSK (15). In myeloid cells, the interaction of Lyp/PEP with TRAF3 promotes TLR activation and type I interferon production (16). Through these studies, researchers have identified mechanisms that may contribute to the pathogenesis of multiple autoimmune disorders. Additionally, we and others found that mice lacking *Ptpn22* (PEP-null) exhibit less of an exhausted T cell phenotype during chronic viral infection (17, 18) suggesting *Ptpn22* contributes to the generation of T cell exhaustion. Furthermore, Brownlie et al have demonstrated adoptively transferred OT-1 CD8 T cells lacking *Ptpn22* have increased function against tumor associated antigens (19, 20). These studies (including our own) were performed using PEP-null mice rather than in mice expressing the equivalent of the human pro-autoimmune allele, it remained to be determined whether the human allele would affect disease pathogenesis in a non-autoimmune context.

Although some studies have found that lack of *Ptpn22* is similar in phenotype to the R620W mutation (R619W in mice) there are also a number of differences. This may be due to the fact that the substitution of arginine to tryptophan within the C-terminus region of the protein does not directly affect phosphatase activity of this enzyme (21). Rather, the presence of tryptophan disrupts the ability of the enzyme to bind with other proteins such as CSK and TRAF, which help direct the phosphatase to appropriate substrates (22, 23). As an example of differing consequences, whereas PEP-null mice exhibit an increased frequency and potency of Tregs as compared with WT (24), this same Treg phenotype is not observed in naïve mice bearing the *Ptpn22* pro-autoimmune allele (1). However, it has been reported that under lymphopenic conditions, both PEP-null and pro-autoimmune allele expressing CD4 T cells have a proliferative advantage over WT CD4 T cells and have a higher propensity to differentiate into Tregs (25). Despite its importance to human health, the *Ptpn22* alternative allele has not been studied in the context of the anti-tumor response, and the accompanying T cell activation and exhaustion. In this study, we employ the use of C57BL/6 mice mutated using CRISPR/Cas9 to express the murine ortholog of the *Ptpn22* pro-autoimmune allele (PEP-619WW), and well-established models of syngeneic murine tumors to define the effect this allele has tumor growth and the anti-tumor response.

## Results

### Effects of the Ptpn22 Pro-autoimmune Allele on Tumor Growth Kinetics

To determine if the human relevant pro-autoimmune allele of *Ptpn22* (PEP-619WW) influenced tumor control, we compared the growth of an immunogenic tumor cell line, B16-OVA, between PEP-WT and PEP-619WW mice. B16-OVA tumors were significantly smaller in PEP-619WW mice than PEP-WT mice one month post implant (Figure 1A-C). The role of the immune system in combating tumor growth is well established (26). Using Rag1^-/-^ mice, which lack T and B lymphocytes, crossed to our PEP-619WW mice (Rag1^-/-^ PEP-619WW mice, Supp. Figure 1A), we tested the necessity of adaptive immunity to control tumors in our mice bearing the *Ptpn22* pro-autoimmune allele. Rag1^-/-^ PEP-619WW mice showed no detectable difference in B16-OVA tumor volume compared to Rag1^-/-^ PEP-WT mice (Supp. Figure 1B-D), suggesting the 619WW mutation promoted an adaptive immune response to tumor.

**Figure 1.**
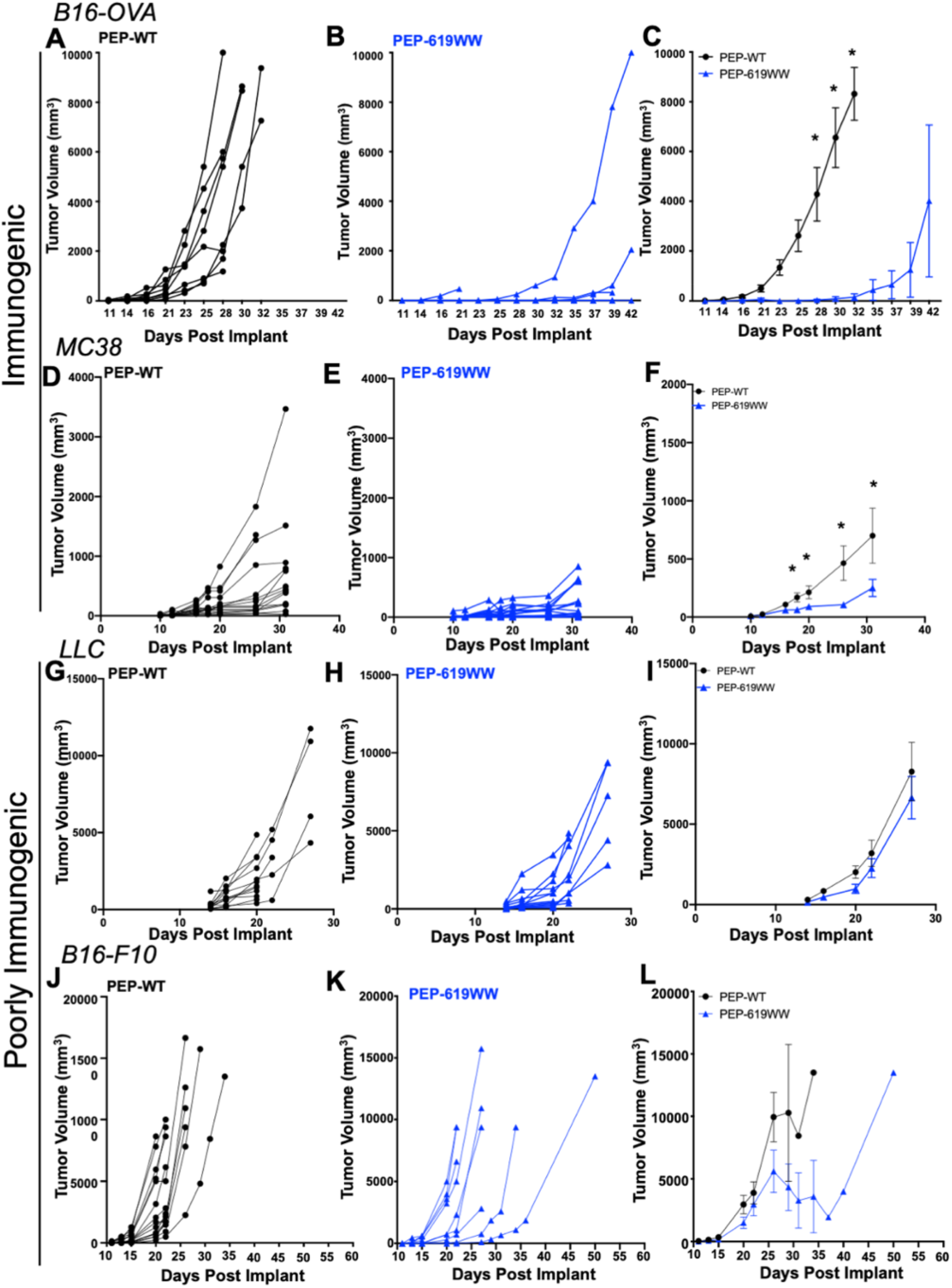
*Ptpn22* pro-autoimmune allele bearing mice have less antigenic tumor burden. PEP-WT (A,D,G,J) or PEP-619WW (B,E,H,K) mice were implanted (sub cutaneous) with 2×10^5^ B16-OVA cells (n=8 each genotype) (A-C), 1×10^6^ MC38 cells (n=23 -WT; 24 -619WW) (D-E), 5×10^5^ LLC cells (n=14 -WT; 13 -619WW) (G-I) or 2×10^5^ B16F10 (n=15 -WT; 14 -619WW) (J-L). Tumor width and length were measured, and volume was calculated. Individual mice for each tumor type shown in A-B, D-E, G-H, J-K. Mean with SEM shown in C,F,I, L. Significance was determined using a T Test with Welch’s correction at each time point to compare the two genotypes; *p<0.05.

Additionally, we tested growth of MC38, another immunogenic tumor line, and found growth was also significantly delayed in PEP-619WW mice (Figure 1 D-F). In contrast, there was no detectable difference in growth of Lewis Lung Carcinoma cells (LLC) between PEP-WT and PEP-619WW mice (Figure 1 G-I). Also, when implanted with B16F10 cells, which are relatively poorly immunogenic, tumor growth was highly variable in PEP-619WW mice, whereas growth in PEP-WT mice was more consistent (Figure 1 J-K). About 1/3 of the PEP-619WW mice had delayed growth, and one mouse which did not develop a palpable tumor until almost a month post implant (Figure 1K). Despite these few mice that seemed to have superior control of B16F10 tumors, as a group there is no significant difference between B16F10 tumor growth in PEP-WT and PEP-619WW mice (Figure 1L). Taken together these results suggest the expression of PEP-619WW can inhibit the growth of immunogenic tumors.

### Ptpn22 pro-autoimmune allele influences Tumor Infiltrating Immune Cells

With PEP-619WW mice having increased tumor control of B16-OVA and MC38 tumors, we set out to define any changes within the immune microenvironment which may be contributing to this delayed tumor growth. We isolated tumor infiltrating lymphocytes (TILs) at 14 days post B16-OVA tumor implant. At this time point, tumors are still small, but there are sizeable numbers of TILs which are responsible for immune control of the tumor. Additionally, this time point is when the size of tumors is comparable in PEP-WT and -619WW mice and would provide insight into immune changes occurring just prior to divergence in growth kinetics. Tumors from PEP-619WW mice were found to have a high frequency of tumor infiltrating CD45+ cells (Figure 2 A-B) at day 14. We also examined the density of CD45+ cells at a later timepoint when B16-OVA tumors are different sizes, day 30. The density of CD45+ tumors was highly variable in both genotypes. There was a trend of increased CD45+ density in B16-OVA tumors from PEP-619WW at both timepoints (Figure 1C). In preparing for an in-depth histochemical analysis of immune rich regions in the tumor, tumor sections from PEP-WT and PEP-619WW mice were stained with anti-mouse CD45. This staining confirmed the flow cytometry data, allowing visualization of CD45+ infiltrate in the tumors from PEP-619WW mice (Figure 2E). Tumors from PEP-619WW mice have more CD45+ cells throughout the tumor. Additionally, the CD45 staining appeared to be more uniform in tumors from PEP-619WW mice, whereas CD45+ staining in tumors from PEP-WT mice was only in distinctive regions in the tissue (Figure 2E). CD45+ tumor infiltration is more robust in mice bearing the *Ptpn22* pro-autoimmune allele.

**Figure 2.**
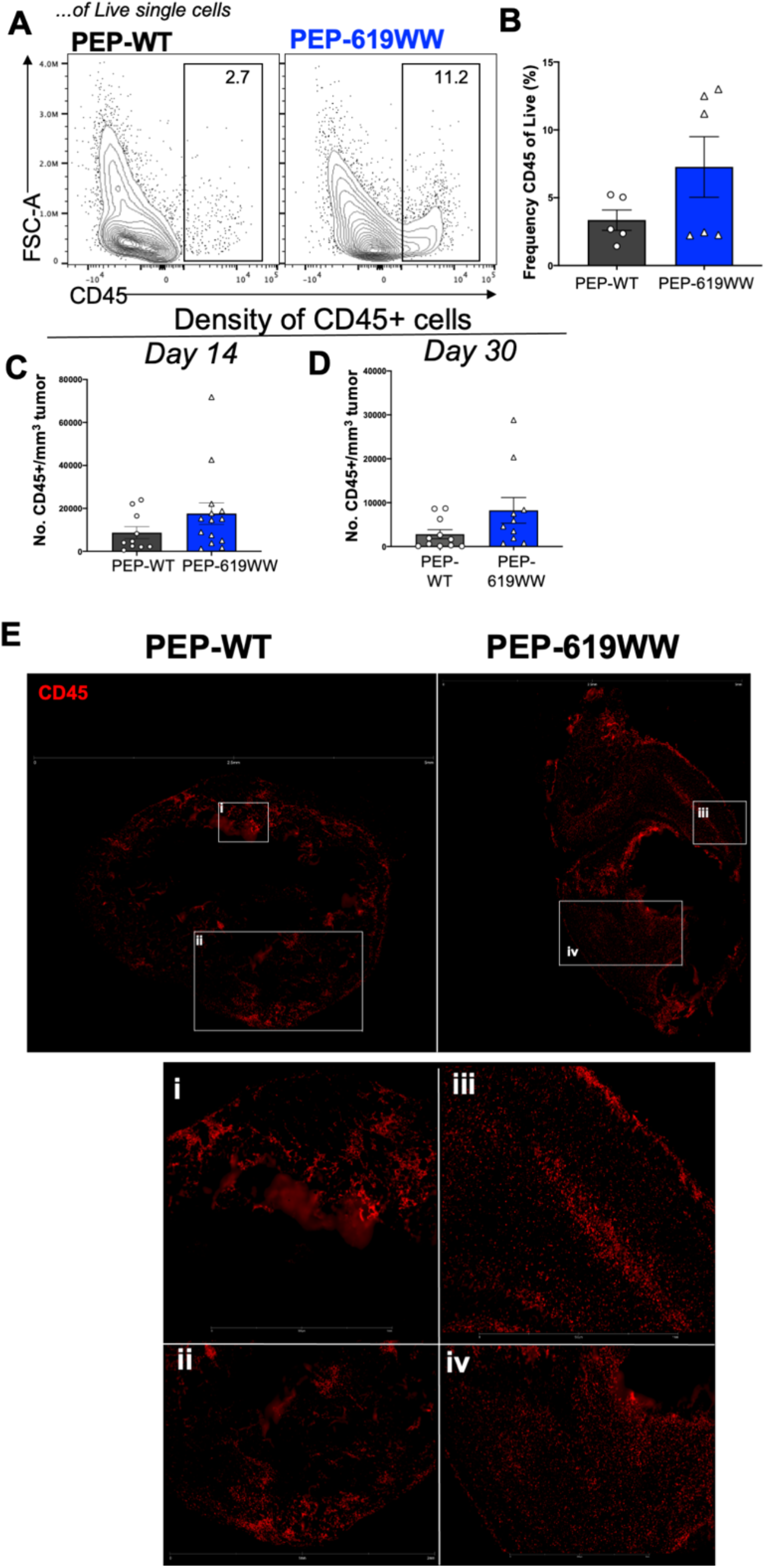
Tumors from PEP-619WW mice have more immune cell infiltrate. B16-OVA tumor infiltrating CD45+ cells were identified by flow cytometry. A) Representative flow cytometry plots (Lymphocytes>Single cell FSC> Single cells SSC> Live> CD45+) and B) quantification of CD45+ tumor infiltrate, determined by flow cytometry. Density of CD45 cells in tumor (count total CD45/volume post resection tumor) at C) 14 days and D) 30 days post tumor implant. E) Representative microscopic images from Nanostring GeoMX DSP platform indicating regions of interest within these shown tumor sections showing CD45 stain (red). Zoomed in images of representative regions of tumors from PEP-WT (i-ii) and PEP-619WW (iii-iv).

### Tumors from PEP-619WW mice have more activated T cell populations

Considering the correlation between increased immune infiltration in tumors from PEP-619WW mice and reduced tumor growth, it was important to define the infiltrating immune cell types in within the tumor infiltrating milieu. We more closely examined T cell activation and myeloid cell activation. Using the Nanostring Spatial Genomic GeoMX Digital Spatial Profiling (DSP) platform, we were able to conduct an in-depth profile of immune related proteins within the immune dense regions of the tumor (day 14 post implant), which were identified with anti-mouse CD45-Alexa532 stain overlaying with DNA-SYTO13. We selected 12 regions of interest in geometrically distinct areas selected over 3 tumor sections for each genotype, where each section was from a different tumor (Supp. Figure 2A-B). To define the differences within the T cell compartment, we first focused on the relative expression of T cell related proteins normalized to CD45 expression within each region of interest, which is measured by the signal-to-noise ratio (SNR). This revealed a significant increase in the expressed levels of CD3e, CD4, CD8a, CD44, CD28, CD27, PD-1, GITR, and Lag-3 in the immune rich regions of tumors from PEP-619WW mice (Figure 3A). Additionally, there is a decrease of CTLA-4 abundance within the immune rich regions of tumors from PEP-619WW mice (Figure 3A).

**Figure 3.**
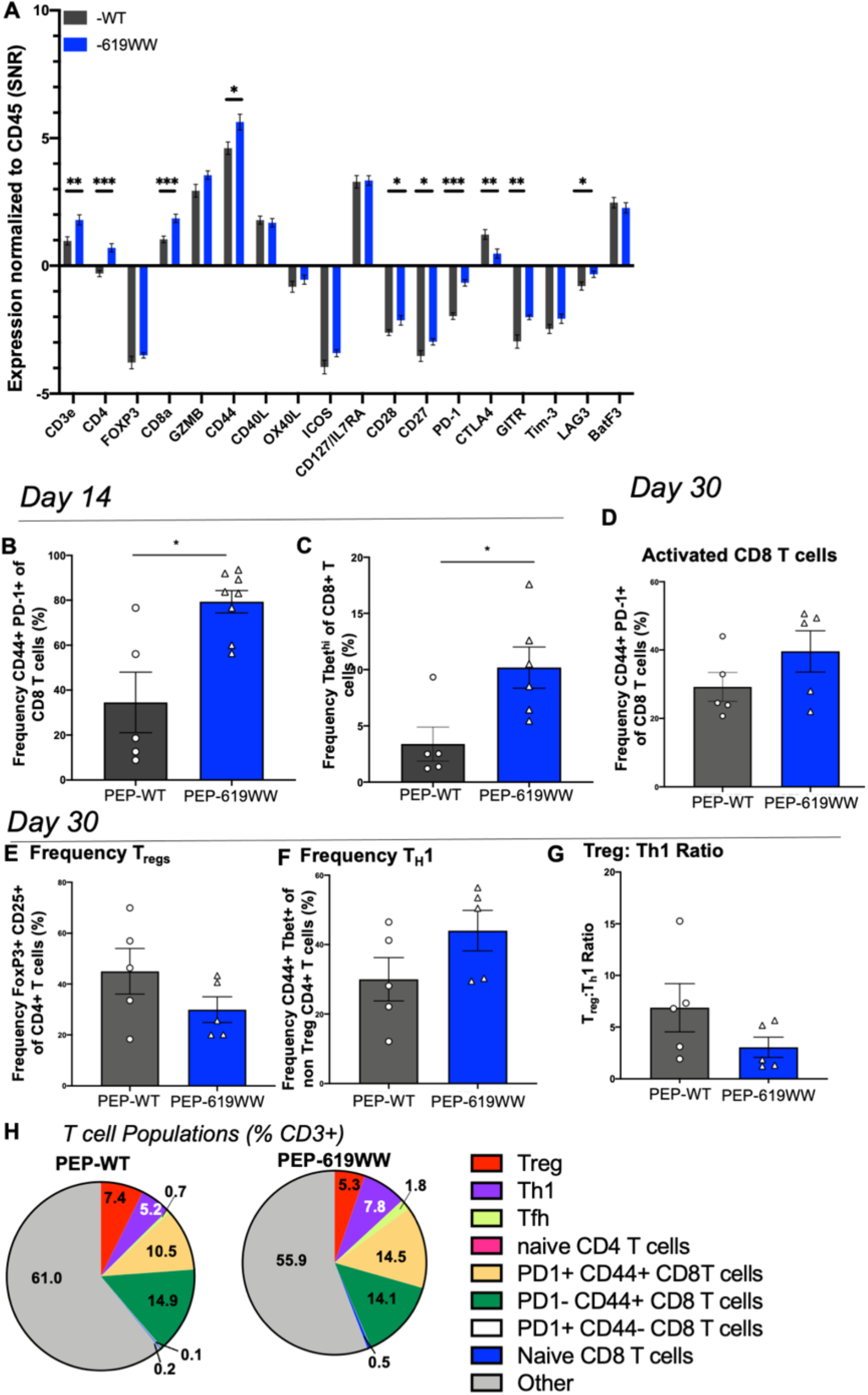
Enhanced T cell activation in tumors from PEP-619WW mice. A) Relative abundance of indicated proteins (normalized to CD45+) within CD45+ rich regions of 14 days post implant B16-OVA tumors, data from Nanostring GeoMX DSP platform. Flow Cytometry analysis of whole tumor at 14 days post implant, B) Frequency CD44+ PD1+ Cd8 T cells (Lymphocyte>Single cellx2> Live> CD45+> CD3+ CD19-> CD8a+ CD4->CD44+ PD-1+), C) Tbet hi CD8 T cell (Lymphocyte>Single cellx2> Live> CD45+> CD3+ CD19-> CD8a+ CD4->Tbethi). E) Frequency activated CD8 T cells 30 days (same gating strategy as B). Frequency of E) Tregs (Lymphocyte>Single cellx2> Live> CD45+> CD3+ CD19-> CD8a-CD4+>FoxP3+ CD25+), F) Th1 (Lymphocyte>Single cellx2> Live> CD45+> CD3+ CD19-> CD8a-CD4+>FoxP3-> CD44+ Tbet+) G) ratio Treg to _TH1_ (%Treg/%T_H_1) and H) breakdown of multiple T cell phenotypes 30 days post implant. T_FH_ cells identified as non Tregs> CXCR5+ PD-1+. T Test with Welch’s correction, *p<0.05

Although this data set allowed us to investigate immune rich regions of the tumors, it did not look at the tumor as a whole, or the level of expression of T cell related markers on specific types of T cells. As such, we next turned to flow cytometry to globally examine T cell activation at a single cell level. Within PEP-619WW mice, the CD8 T cell compartment, exhibited a more activated phenotype as measured by CD44 and Tbet expression at 14 days post implant (Figure 3B-C). The increased T cell activation present at 14 day in tumors from PEP-619WW mice persisted at 1 month post implant. At this later time point, tumor from PEP-619WW retained an increased frequency of activated CD8 T cells (CD44+ PD-1+ CD8 T cells) (Figure 3D,H). Additionally, PEP-619WW had altered CD4 T cell phenotypes present within the tumor as compared to tumors from PEP-WT mice. Tumors from PEP-619WW mice had a decreased frequency of Tregs (FoxP3+ CD25+ CD4 T cells), and increased frequency of T_H_1 cells (non Tregs CD44+ Tbet+ CD4 T cells) (Figure 3E-F,H). This rendered a significantly lower Treg:T_H_1 ratio in tumors from PEP-619WW mice (Figure 3G). Importantly, there were no differences in the number of Tregs in the spleens of naïve animals of either genotype, a phenotype which was previously reported as a characteristic of PEP-null mice (Supp. Figure 3 and (24)). There was also increased proportion of T follicular helper cells (T_FH_) in tumors from PEP-619WW mice compared to PEP-WT mice (Figure 3H).

### PEP-619WW mice have increased myeloid cell activation in tumor microenvironment

Although T cells are necessary for proper tumor control, the dendritic cell and myeloid compartment is also critical in supporting a cytotoxic T cell response (27, 28). Additionally, the presence of suppressive myeloid cells can counteract a strong T cell infiltrating response through expression of T cell inhibitory ligands (like PD-L1) and suppressive cytokine secretion (27). Again, using the Nanostring Spatial genomics GeoMX/DSP platform, we focused on immune dense regions within the tumors from PEP-WT and PEP-619WW mice to broadly examine myeloid and tumor associated markers. Tumors from PEP-619WW mice had significant increase in the abundance of cellular identification markers CD19 and CD11c. Additionally, tumors from PEP-619WW mice had increased amounts of T cell activating receptors CD40 and MHC II within the immune rich regions (Figure 4A). Also, these same areas had a lesser abundance of myeloid derived suppressor cell marker, CD163, and tumor associated receptors B7-H3, and Androgen receptor (AR), which have been previously correlated with poor tumor clearance (Figure 4A, (29-31)).

**Figure 4.**
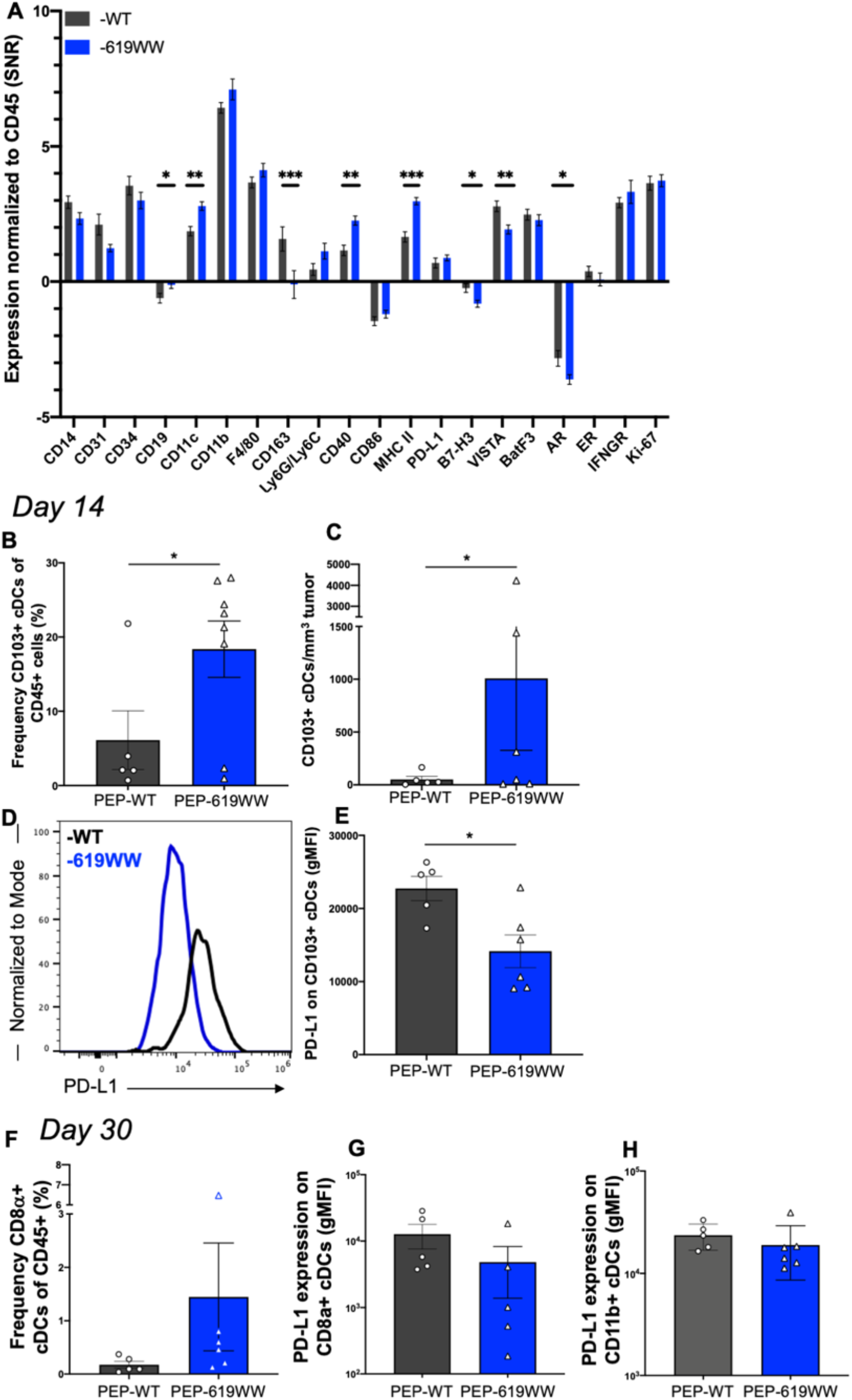
Enhanced myeloid cell activation in tumors from PEP-619WW mice. A) Relative protein abundance (normalized to CD45) of indicated protein target from CD45 rich regions of interest in B16-OVA tumors from PEP-WT and PEP619WW mice, 14 days post implant. Data from Nanostring GeoMX DSP platform. Tumors were harvested at 14 days post implant and put in a single cell suspension for flow cyyometry analysis, B) Frequency of tumor CD103+ cDCs of CD45+ cells (Lymphocytes> Single cellsx2> Live> CD45+> CD19-CD3-> F4/80->CD11c+ MHC II+ >CD103+> CD8a+ CD11b-). C) Density of CD103+ cDCs in tumors at 14 days post implant. D) Representative histogram and E) quantification of PD-L1 surface expression on tumor infiltrating CD103+ cDCs at 14 days post implant. F) Frequency of CD8a+ cDCs of CD45+ tumor infiltrating cells (Lymphocytes> Single cellsx2> Live> CD45+> CD19-CD3-> F4/80->CD11c+ MHC II+> CD8a+ CD11b-), 30 days post implant. PD-L1 surface expression (gMFI) on G) CD8a+ cDCs and H) CD11b+ cDCs (Lymphocytes> Single cellsx2> Live> CD45+> CD19-CD3-> F4/80->CD11c+ MHC II+> CD8a-CD11b+). Significance was determined using a T test with Welch’s correction, *p<0.05, **p<0.01, ***p<0.001.

Next, we turned to flow cytometry to examine the immunostimulatory versus immunoinhibitory phenotype of the myeloid cells within the tumor microenvironment from PEP-WT and PEP-619WW mice at a single cell level. We focused on dendritic cells within the tumor microenvironment as these are critical for proper T cell activation and immune mediated tumor clearance (28). Tumors from PEP-619WW mice had a higher proportion, and density of cDC1s (F4/80-CD11c+ MHC II+ CD103+) (Figure 4B-C) at day 14 post implant. Additionally, these cDC1s had significantly lower levels of PD-L1 expression (Figure 4D-E). At day 30 post implant, tumors harvested from PEP-619WW mice had more cDC1 cells (F4/80-CD11c+ MHC II+ CD8a+ CD11b-) (Figure 4F). At this later timepoint, there was large variability in the level of expression of PD-L1 on cDC1s in the PEP-619WW mice (Figure 4G), whereas PEP-WT tumor infiltrating cDC1 cells had consistently high levels (Figure 4G). Infiltrating cDC2 cells (F4/80-CD11c+ MHCII+ CD8a-CD11b+) had consistently high levels of PD-L1 expression (Figure 4H).

## Discussion

There is a strong genetic link associated with autoimmunity which drives overt immune activation (2, 32). Given this relationship, we wanted to know if expression of a pro-autoimmune allele could drive a positive disease outcome in situations in which overt immune activation is favorable, such as cancer. Previously, using mice deficient in *Ptpn22* expression, we, and others, have demonstrated that the protein tyrosine phosphatase *Ptpn22* drives T cell exhaustion during viral infection (17, 18). Also, Brownlie et al, showed CD8 T cells lacking *Ptpn22* were superior as adoptive cell transfer therapy of tumors compared to wild type CD8 T cells (19, 20). Recently, another group showed *Ptpn22* knock out mice had enhanced rejection of MC38 and Hepa1-6.x1 during immunotherapy with checkpoint blockade (33). This same study briefly looked at the spontaneous rejection of Hepa1-6.x1 cells in mice bearing the *Ptpn22* pro-autoimmune allele (PEP-619WW mice). There was a modest increase in spontaneous rejection compared to WT animals (33). However, the study did not look at other tumor lines nor elaborate on the underlying immunological differences between the wildtype and the mice bearing the pro-autoimmune allelic variant of *Ptpn22*.

To test any influence the *PTPN22* pro-autoimmune allelic variant has on tumor growth and the anti-tumor immune response, we used CRISPR/Cas9 to generate C57BL/6 mice which endogenously express the murine ortholog of the human relevant *Ptpn22* variant (PEP-619WW). We tested the capacity of our PEP-619WW mice to control both immunogenic (B16-OVA and MC38), and poorly immunogenic (B16F10 and LLC) tumors (Figure 1). Following implantation of B16-OVA cells, PEP-619WW mice exhibited significantly better tumor control without any therapeutic intervention (Figure 1 A-C). PEP-619WW mice also had superior control of another immunogenic tumor, MC38 (Figure 1 D-F). However, when PEP-619WW mice were implanted with the less antigenic B16F10 melanoma cells, this significant increase in tumor control was mitigated (Figure 1 J-L). Furthermore, like their WT counterparts, PEP-619WW mice were not able to control the poorly immunogenic LLC tumor (Figure 1 G-I). These data suggest that the enhanced tumor control seen in PEP-619WW mice was driven by the presence of novel tumor expressed antigens.

It is known that T cell activation is critical for the cytotoxic destruction of malignancies and is associated with improved survival of melanoma (34, 35). Indeed, using a Rag1^-/-^ mice, which lacks lymphocytes, expressing PEP-619WW in the innate compartment can no longer control B16-OVA tumors (Supp. Figure 1B-D). In our immune competent mice, tumors from PEP-619WW had more immune cell infiltration at both an early (14 days post implant) and later (30 days post implant) timepoints (Figure 2). When we focused on the immune rich regions of the tumors (at day 14) using the Nanostring GeoMX DSP platform, there was increased relative expression of CD3 and CD8a, suggesting more CD8 T cell infiltrating in tumors from PEP-619WW mice (Figure 3A). Furthermore, there was more CD44 and PD-1 expression, which are upregulated upon T cell activation, in tumors from PEP-619WW mice (Figure 3A). Lastly, there was less CTLA-4 and more CD28 relative expression in tumors from PEP-619WW compared to PEP-WT (Figure 3A). CTLA-4 binds to CD80 restricting further activation through the interaction of CD80 and CD28 (36, 37). With lower levels of CTLA-4 and increased levels of CD28 within the tumors from PEP-619WW suggests that these T cells are receiving more activating than suppressive signals. Using flow cytometry, we were able to assess T cell activation on single cell level and confirmed that tumors from PEP-619WW mice had significantly more activated CD8 T cells infiltrating at 14 days post implant (Figure 3C-D). We also show that tumors from PEP-619WW have a persistent CD8 T cell activation present at 30 days post tumor implant (Figure 3E). It is well characterized that the pro-autoimmune allele of *Ptpn22* can enhance CD8 T cell activation (38), to our knowledge this is the first such demonstration within the tumor microenvironment. Future research will assess the killing capacity of the tumor infiltrating CD8 T cells from PEP-619WW and if expressing this pro-autoimmune allele has any advantage for adoptive cell therapy, as seen for T cells lacking *Ptpn22* does (19, 20).

CD4 T cell subsets also play a key role in the anti-tumor response (39, 40). Previous studies have shown that *Ptpn22*, and its pro-autoimmune allele, do influence CD4 T cells activation and function (41). T regulatory cells (Tregs) have are associated with poor tumor control, as these cells can temper cytotoxic T cell function (37). Mice lacking *Ptpn22* have more Tregs and increased potency of Treg function compared to wildtype mice (24, 42). However, mice bearing the pro-autoimmune allele of *Ptpn22* do not have an increased amount of Tregs under homeostatic conditions ((1), Supp. Fig 3). In this study we show that at one month post implant, PEP-619WW actually have fewer tumor infiltrating Treg compared to tumors from PEP-WT mice (Figure 3F, H-I). Additionally, tumors from PEP-619WW mice had more T_H_1 and T_FH_ cells, both of which are reported to correlate with positive outcomes in cancer, compared to tumors from PEP-WT mice ((40), Figure 3G-I). Taken together, these data suggest that the *Ptpn22* pro-autoimmune allele promotes a more activated tumor infiltrating T cell population. Another important consideration for proper tumor control is the presence of myeloid derived suppressor cells (MDSCs). *Ptpn22* can influence the polarization of macrophages (43, 44). Within immune rich regions, we did observe that PEP-WT mice had more CD163 relative expression, suggesting a higher frequency of MDSCs present in the tumor (Figure 4A). Although tumors from PEP-619WW mice had high level of CD11b expression, they did not have high levels of CD163 expression, suggesting these were not MDSCs in the tumor microenvironment (Figure 4A).

Within these immune rich regions, we also observe increased expression of CD11c, suggesting more dendritic cells (DCs), in tumors from the PEP-619WW mice (Figure 4A). Tumors from PEP-619WW mice also exhibited greater expression levels of CD40 and MHC II, both which serve as activators of the T cell response (Figure 4A). Additionally, there is less inhibitory checkpoint receptor VISTA expression in tumors from PEP-619WW mice ((45) Figure 4A). Taken together, this suggests that the tumor infiltrating DCs in PEP-619WW are better equipped for T cell activation.

To interrogate the DCs at a single cell level, we again turned to flow cytometry. Confirming the GeoMX DSP data, there were more tumor infiltrating DCs in PEP-619WW, specifically conventional DC type 1 (cDC1) cells (Figure 4B-C,F). cDC1 cells are critical for priming a CD8 T cell response and mounting a robust and proper anti-tumor immune response which leads to better tumor control (28). More cDC1s has been associated with better tumor control and a more activated CD8 T cell phenotype. Not only were the cDC1s more abundant in the PEP-619WW tumor bearing mice, these cells also had significantly less inhibitory receptor PD-L1 surface expression (Figure 4D-E). To our knowledge, we are the first to put forward a link between this pro-autoimmune allele and PD-L1 expression. The difference in PD-L1 expression between PEP-619WW and PEP-WT cells was specific to the cDC1 compartment, cDC2 cells (CD11b+ CD8a-CD11c+ MHC II+) did not exhibit a difference in PD-L1 expression within the tumor (Figure 4H). Clinically, the blockade of PD-L1 and is cognate receptor PD-1 has had success in promoting tumor clearance (46, 47). In the B16-OVA system, tumor control is greater when anti-PD-L1 is paired with an antigen loaded DC immunization and OVA-specific T cells adoptive transfer therapy (48). Future studies will define the molecular mechanism the PEP-619WW molecule is affecting PD-L1 expression and further define if this difference in PD-L1 expression between the two genotypes is required for tumor control.

This study explores the role of the tyrosine phosphatase *PTPN22* pro-autoimmune allele on tumor control. Broadly, we have determined that pro-autoimmune alleles can influence the anti-tumor response by promoting a more pro-inflammatory environment and overcoming the anti-inflammatory signals present during tumor growth. Interestingly, the presence of this pro-autoimmune allele affects both the T cell and non-T cell compartment to promote better control of tumor growth. In humans the *PTPN22* pro-autoimmune allele, rs2476601, is associated with a lower incidence of melanoma and other malignant neoplasm of the skin (C43-C44) (p=1.8196e-8; Gene Atlas UK biobank population). Currently, we do not know if the enhancement in the T cell response, myeloid cell activation, or both is driving the difference in tumor control between our 2 strains of mice. Transfer of PEP-619WW T cells into a lymphopenic environment (such as into a Rag1^-/-^ mouse or lethally irradiated animal) preferentially differentiate into Tregs at a higher proportion compared to PEP-WT T cells (25). Therefore, isolating lymphocytes or myeloid cells as the sole *Ptpn22* differing cell through adoptive transfer techniques into immune deficient hosts do not properly recapitulate the phenotypes observed in our immune competent mice. Future studies will need to examine which PEP-619WW bearing immune cells are necessary and/or sufficient to control tumors as well as understand if these results could translate into improved therapies for patients. Results of this study provide a platform to further investigate the concerted relationships between immune cells to achieve a desired phenotype as well as determine if/how other pro-autoimmune alleles affect the anti-tumor immune response.

## Methods

### Mice

Both males and females ranging from 6-12 weeks of age were used in this study. Animals were housed in general housing conditions at Scripps Research-La Jolla. All animal studies were reviewed and approved by Scripps Research Institutional Animal Care and Use Committee (protocol number: 06-0291). C57BL/6 WT mice were originally purchased from Jackson Labs, and then bred and maintained in Scripps Animal Facility. *Ptpn22* pro-autoimmune allele bearing mice (PEP-619WW) were generated using CRISPR/Cas9 technology on a C57BL/6 background using methods previously reported (5). In short, four nucleotides were replaced on exon 14 of *Ptpn22* to insert the BspEI restriction site and cause an arginine (R) to tryptophan (W) amino acid substitution at amino acid position 619. Genotypes were confirmed through PCR using the following primers which flank the mutated region of *Ptpn22*: Forward-5’ AGCTGATGAAAATGTCCTATTGTGA 3’ and Reverse-5’ GTCCCACTGCATTCTGGTGA 3’. After amplification, PCR products are digested overnight at 37°C with the restriction enzyme, BspEI, which is unique to mutated mice. Digested PCR products are run on an agarose gel to visualize digested bands (Supplemental Figure 4).

Rag1^-/-^ C57BL/6 mice were a gift from Dr. David Nemazee (Scripps Research-La Jolla) and then were maintained in our colony. These mice were crossed with our PEP-619WW C57BL/6 mice to created Rag1^+/-^ PEP-WT/619W heterozygotes. These mice were backcrossed to Rag1^-/-^ PEP-WT to generate Rag1^-/-^ PEP-WT/619W heterozygous mice. These mice were bred to each other to eventually generate mice which lacked lymphocytes and were homozygous for the pro-autoimmune allele of *Ptpn22*, which we denote as Rag1^-/-^ PEP-619WW mice (Breeding scheme diagramed Supp. Figure 1). These mice were genotyped by lack of circulating lymphocytes and with the PCR genotyping protocol described above.

### Tumor Culture and Growth

All tumor cell lines were cultured in Dulbecco’s Modified Eagles Medium (DMEM) supplemented with 10% heat inactivated fetal calf serum (FCS), 1% glutamine and gentamicin, and 50mM β-mercaptoethanol at 37°C, 5% CO_2_. Additionally, B16-OVA cells were cultured with geneticin (G418) for at least 2 passes before implanting in mice. Cell lines are passed once they are at 80% confluency. Cells are not cultured for greater than 3 months.

One day prior to tumor implant, the left flank of mice are shaved. While tumor cells are in growth phase, based on confluency of plate (less that 80% confluent), cells are harvested, counted, and resuspending in sterile HBSS for implantation into mice. Tumor cells implanted subcutaneously in shaven area of mice at the following amounts (all in a total volume of 100uL): 2×10^5^ B16-OVA, 1×10^6^ MC38, 2×10^5^ LLC, or 2×10^5^ B16F10 cells. Mice were monitored and tumor growth was measured at indicated time points. Tumor volume (mm^3^) was calculated with the following equation: [Length x width^2^]/2. Once the tumor reached 30mm in any direction, or became ulcerated, mouse was removed from the study.

### Flow Cytometry and Antibodies

Tumors were excised and placed into HBSS with 2% FBS. Samples were minced and incubated with Stem Cell Spleen dissociation media (Stem Cell Technologies, Vancouver, British Columbia, Canada) according to manufacturer’s instructions. Following incubation, minced sample was smashed and filtered through 40uM filter to create a single cell suspension. Single cell suspension was counted and resuspended to desired concentration (dependent on experiment) in HBSS with 2% FBS. Single cell suspensions were used for staining and flow cytometric analysis. Cells were stained in serum free HBSS.

All flow cytometry was completed on a spectral cytometer the Cytek Aurora with a 4 laser or 5 laser system (405nm, 488nm, 640nm, 561nm, and 355nm (5 laser only)). Single color stain OneComp eBeads (Thermo Fisher, United States) were used for unmixing. Unmixed files were analyzed using FlowJo Software (BD Biosciences, San Diego, California). Antibodies used in various combinations (depending on experiment) are as follows: Ghost Viability Dye (v510, Tonbo Biosciences, 1:1000 dilution), CD45 (BV570/BV605, Biolegend, 1:200, clone 30-F11), CD3e (PE-Cy5/AF532, Tonbo Bioscience/Thermo Fischer, 1:200, clone 145-2C11), CD4 (PerCP/BV605, Tonbo Bioscience/Biolegend, 1:200, clone RM4-5), CD8a (APC-Cy7/APC-H7/APC, BD Biosciences, 1:200, clone 53-6.7), CD11c (PE-Cy5.5, Thermo Fisher, 1:100, clone N418), CD11b (PerCP-Cy5.5, Biolegend, 1:200, clone M1/70), F4/80 (Pacific Orange, Thermo Fisher, 1:100, clone BM8), PDCA-1 (Pacific Blue, Biolegend, 1:200, clone 129C1), CD80 (BV421, 1:200, clone 16-10A1), CD86 (BV605, Biolegend, 1:200, clone GL1), PD-1 (PE-Cy7, Tonbo Biosciences, 1:200, clone J43.1), PD-L1 (PE/BV711/PE-Cy7, Tonbo Biosciences/Biolegend, 1:100, clone 10F.9G2), CD44 (AF700, Biolegend, 1:200, clone IM7), CD62L (FITC, Tonbo Biosciences, 1:100, MEL-14), Ly6C (BV785, Biolegend, 1:200, clone HK1.4), Ly6G (PE-eFlour610, Invitrogen, 1:200, clone IA8), CD206 (AF647, Biolegend, 1:100, clone CO68C2), CD209b (APC, Tonbo Biosciences, 1:200, clone 22D1), NK1.1 (FITC, Biolegend, 1:100, clone PK136), CD19 (BV711, Biolegend, 1:400, clone 6D5), B220 (APC-Cy5.5, Invitrogen, 1:200, clone RA3-6B2), MHC II I-Ab (FITC, Biolegend, 1:200, clone AF6-120.1), CXCR5 (BV605, Biolegend, 1:100, clone L138D7), Tbet (APC/AF647, Biolegend, 1:200), and FoxP3 (PE, Invitrogen, 1:100, clone FJK-16s). Cells were stained in HBSS, at 4C, in dark. If intracellular staining for transcription factors was required, Tonbo Bioscience FoxP3 Fix/Perm kit was used per manufacturer’s instructions (Tonbo Bioscience, San Diego, California)

Density of immune cells in tumors was determined by the following equation: [total count CD45+ cells in tumor/tumor volume post resection (mm^3^)].

### Nanostring Spatial GeoMX Digital Spatial Profiling (DSP)

At 14 days post implant of B16-OVA tumors, tumors from PEP-WT and PEP-619WW mice were harvested, fixed in 10% Neutral Buffered Formalin, and paraffin embedded. The 3 mean tumor sizes from each genotype were cut and put onto slides and sent to Nanostring, Seattle, WA for GeoMX digital spatial profiling (GeoMX DSP). Tissue sections were stained for morphology markers DNA-SYTO13), Pmel/S100b-Alexa532, and CD45-Alexa594, and a set of 53 photocleavable probes. Using these morphology markers a total of 12 regions of interest per genotype were selected based on CD45 rich regions spaced throughout each tissue section. Each region of interest was exposed to UV illumination and the eluent was collected and transferred into individual wells of a microtiter plate. Once the 12 ROIs were processed, indexing oligos were hybridized to NanoString optical barcodes for digital counting on the nCouter^®^. Digital counts from barcodes corresponding to protein probes were then normalized to housekeeping counts, area, and CD45 probe. Differences in protein expression (measured by signal to noise ratio-SNR) between genotypes was determined by T Test with Welch’s correction.

List of target proteins: Aryl hormone receptor (AhR), Androgen receptor (AR), B7-H3, Batf3, CD11b, CD11c, CD227/IL17RA, CD14. CD163. CD19, CD27, CD28, CD31, CD34, CD3e, CD4, CD40, CD40L, CD44, CD45, CD86, CD8a, CTLA-4, Epcam, ER (estrogen receptor), F4/80, Fibronectin, FoxP3, GAPDH, GFP, GITR, Granzyme B (GzmB), Her2, Histone H3, ICOS, IFNgR, Ki67, Lag3, Ly6G/C, MHC II, OX40L, PanCK, PD-1, PD-L1, Pmel17, RbIgG, RbIgG2a, RbIgG2b, S100b, S6, SMA, Tim-3, and VISTA.

### Statistics

Statistical tests were completed in Graphpad Prism Software. Specific tests and p value which is determined significant are stated in figure legends.

## Author Contributions

R.C.O. conceptualized, designed and performed experiments, analyzed results, and wrote and edited the manuscript. K.Ma. provided reagents, bred and genotyped mice, and performed experiments. K.Mo. was the principal investigator whose lab generated the PEP-619WW mice on the C57BL/6 background. Since their creation, K.Mo. has passed away and is greatly missed. L.A.S conceptualized and designed experiments, reviewed data, and wrote and edited the manuscript.

## Acknowledgements

We would like to thank Scripps Research Flow Cytometry Core, Histology and Microscopy Core, and Vivarium staff for their expertise and assistance in this work. We would also like to thank C. Alvarez for his help with cutting tissue section as well as careful reading and editing of this manuscript and L. Zhang, Nanostring, for his patience and assistance in going through the data from the Nanostring GeoMX DSP platform. This work was funded by NIH U01 AI130842 (awarded to L.A.S) and NIH T32 AI007354 27 (fellowship to support R.C.O).

---

**Supplemental Figure 1.**
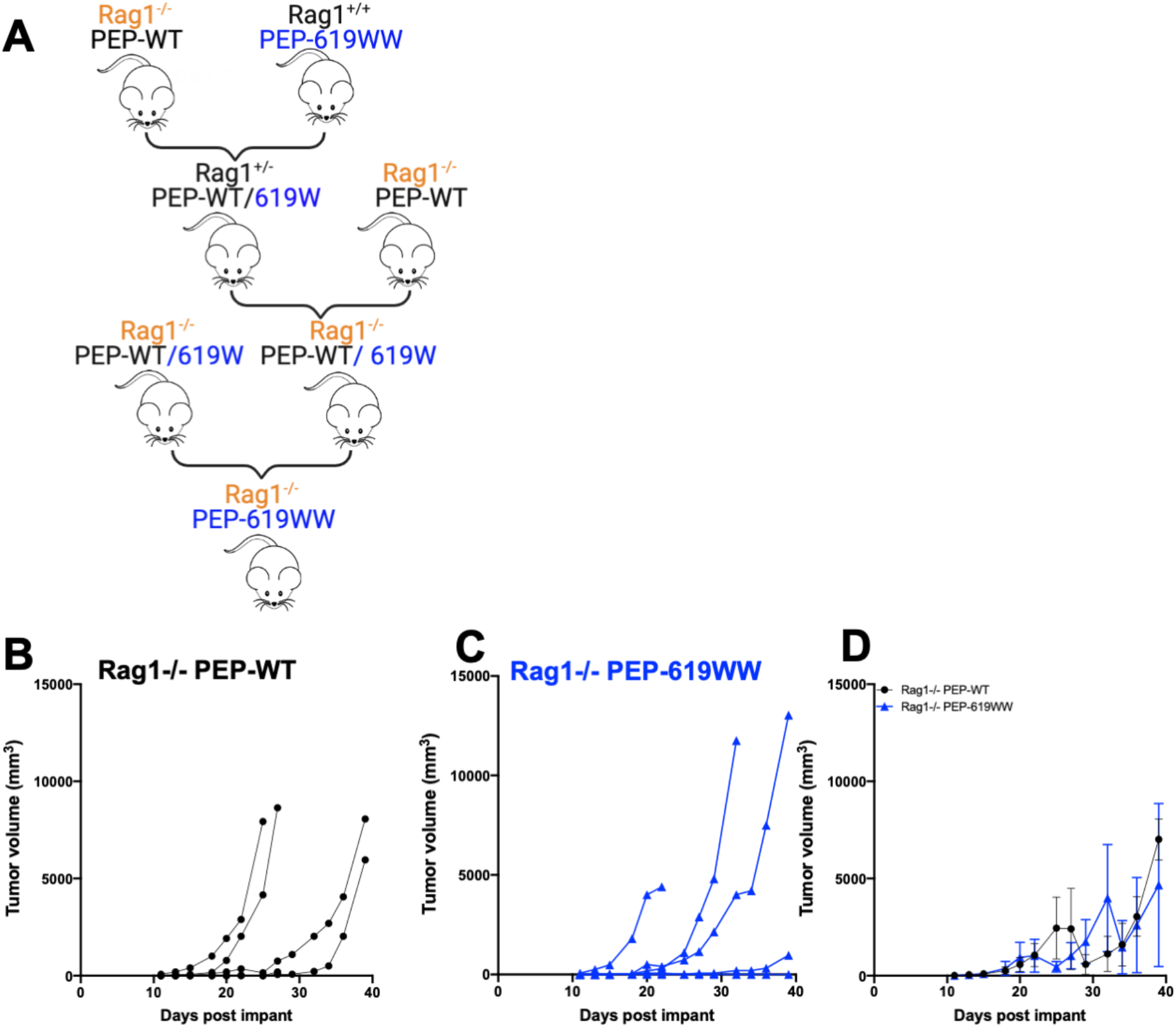
Lymphocytes are needed to control tumors in both PEP-WT and PEP-619WW mice. A) Breeding schematic used to create Rag1^-/-^ mice with PEP-619WW expression. B16-OVA growth curve in B) Rag 1^-/-^ PEP-WT individual mice, C) Rag 1^-/-^ PEP-619WW individual mice. D) Average growth with SEM shown. N=5 for each genotype. T test with Welch’s Correction at each time point was done to determine any significant differences between genotypes.

**Supplemental Figure 2.**
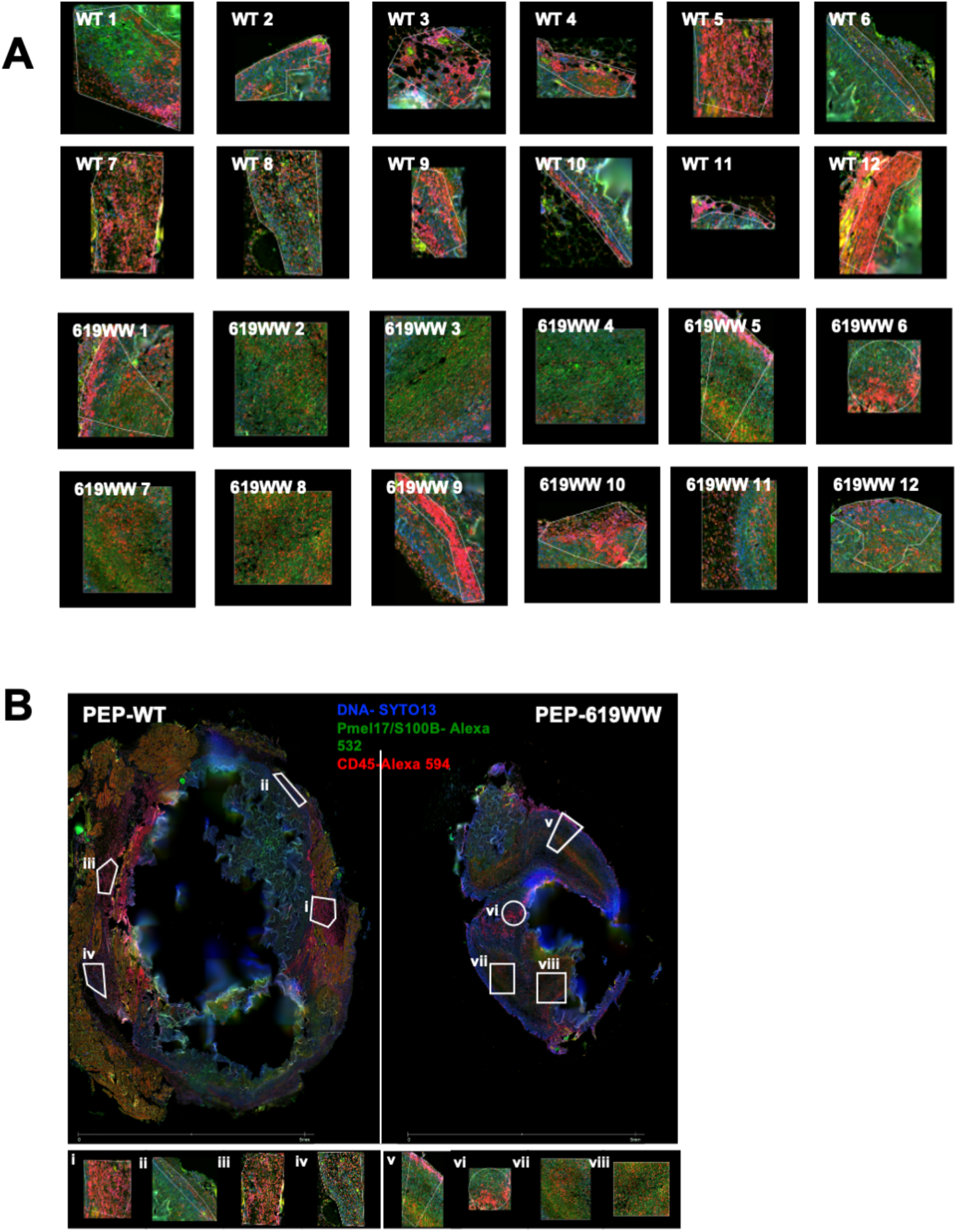
Nanostring GeoMX DSP Selected Regions of interest at CD45 dense regions. A) Individual regions for each genotype (12 for PEP-WT and 12 for PEP-619WW mice). B) Example of 1/3 tumor sections for each genotype with regions of interest outlined in white. N=3 tumors per genotype.

**Supplemental Figure 3.**
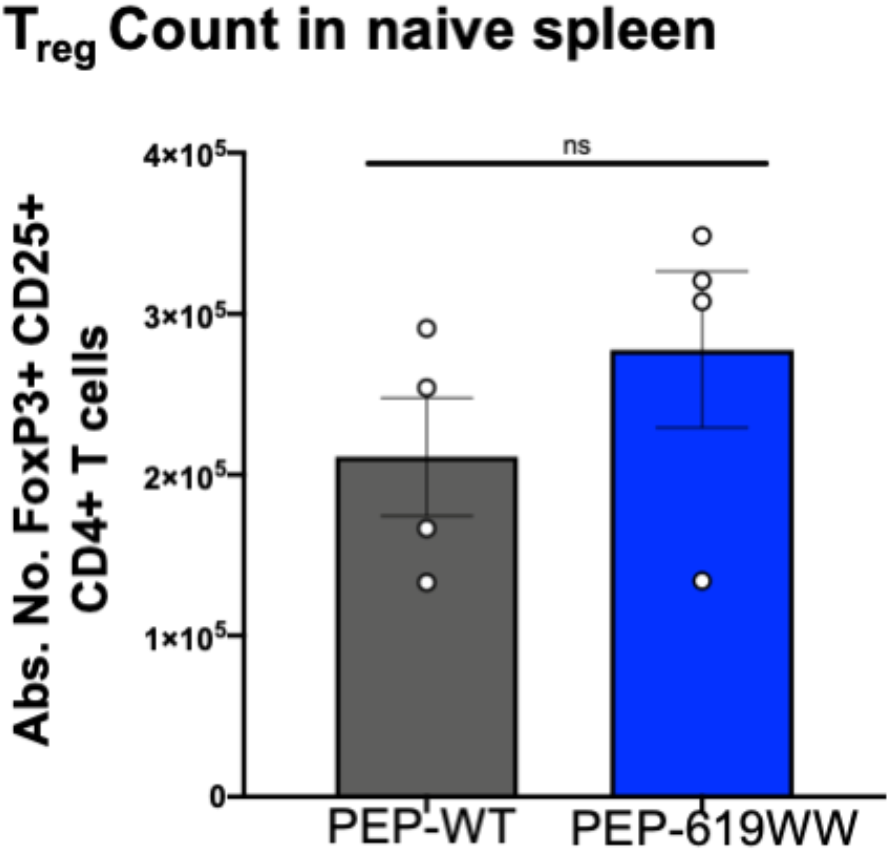
Non detectable difference in splenic Treg count between WT and PEP-619WW mice. Spleens were harvested from naïve PEP-WT and PEP619WW mice. Absolute number Tregs in spleen with SEM shown. Gating strategy: Lymphocyte>Single cellx2> Live> CD45+> CD3+ CD19-> CD8a-CD4+>FoxP3+ CD25+. Significance determined via T Test with Welch’s correction.

**Supplemental Figure 4.**
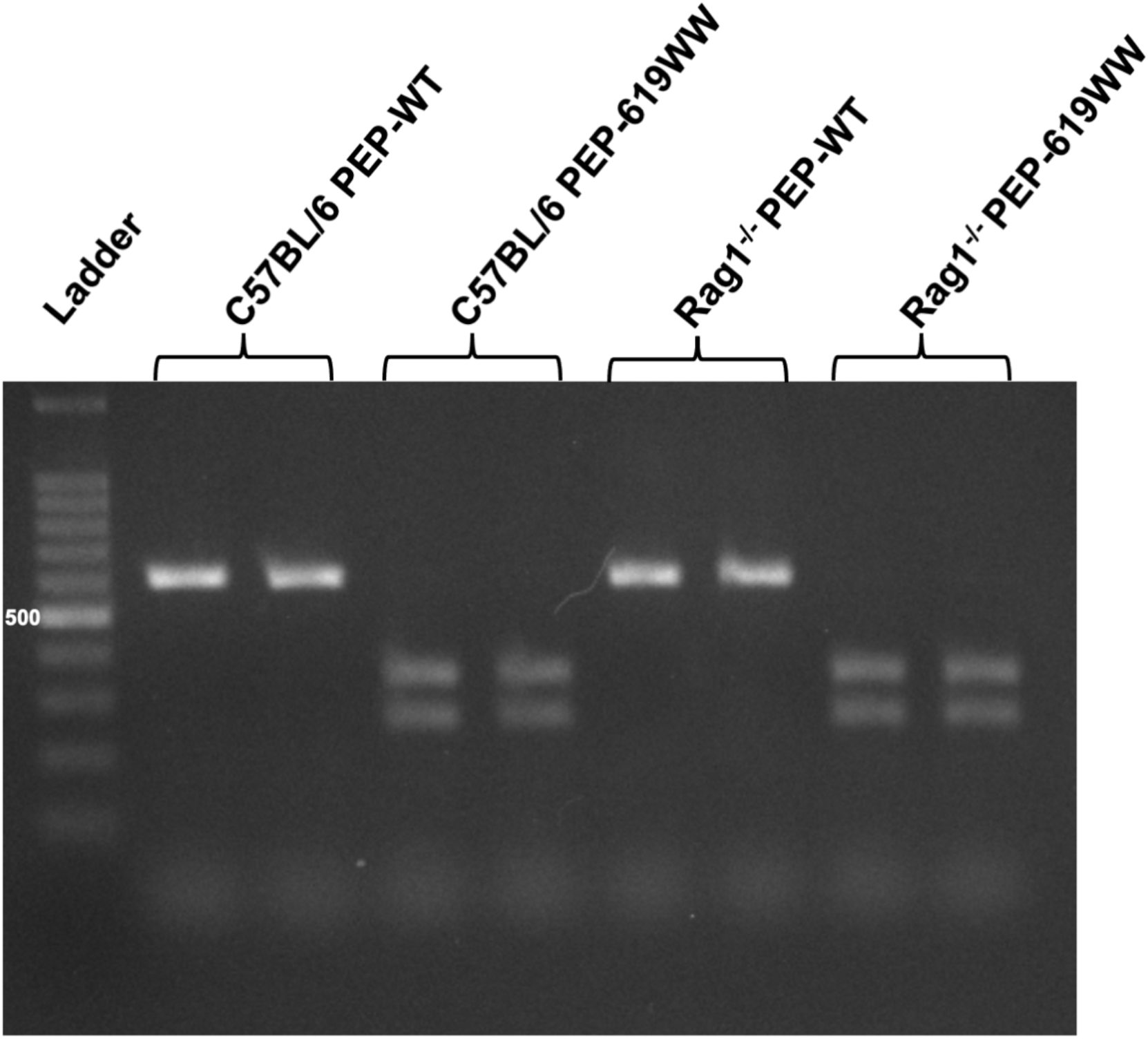
Genotyping PEP-619WW mice. Amplified region of *Ptpn22*, which encompasses mutation sites responsible for the Arg→ Trp amino acid substitution at position 619, is digested with restriction enzyme BspEI. This site is only present in mice with the mutation. Digested fragments are run on an agarose gel and visualized. PEP-WT mice have a single band greater than 500 bp. PEP-619WW mice have 2 band less than 500bp each.

## References

1. X. Dai et al., A disease-associated PTPN22 variant promotes systemic autoimmunity in murine models. J Clin Invest 123, 2024–2036 (2013).

2. G. L. Burn, L. Svensson, C. Sanchez-Blanco, M. Saini, A. P. Cope, Why is PTPN22 a good candidate susceptibility gene for autoimmune disease? FEBS letters 585, 3689–3698 (2011).

3. N. Bottini, T. Vang, F. Cucca, T. Mustelin, Role of PTPN22 in type 1 diabetes and other autoimmune diseases. Seminars in immunology 18, 207–213 (2006).

4. S. Cohen, H. Dadi, E. Shaoul, N. Sharfe, C. M. Roifman, Cloning and characterization of a lymphoid-specific, inducible human protein tyrosine phosphatase, Lyp. Blood 93, 2013–2024 (1999).

5. X. Lin et al., CRISPR-Cas9–Mediated Modification of the NOD Mouse Genome With Ptpn22 R619W Mutation Increases Autoimmune Diabetes. Diabetes 65, 2134–2138 (2016).

6. L. M. Gomez et al., PTPN22 C1858T polymorphism in Colombian patients with autoimmune diseases. Genes and immunity 6, 628–631 (2005).

7. Y. Di, S. Zhong, L. Wu, Y. Li, N. Sun, The Association between PTPN22 Genetic Polymorphism and Juvenile Idiopathic Arthritis (JIA) Susceptibility: An Updated Meta-Analysis. Iranian journal of public health 44, 1169–1175 (2015).

8. R. C. Chiaroni-Clarke et al., The association of PTPN22 rs2476601 with juvenile idiopathic arthritis is specific to females. Genes and immunity 16, 495–498 (2015).

9. Y. Cao et al., PTPN22 R620W polymorphism and ANCA disease risk in white populations: a metaanalysis. The Journal of rheumatology 42, 292–299 (2015).

10. X. H. Wang et al., Protein tyrosine phosphatase nonreceptor type 22 (PTPN22) gene single nucleotide polymorphisms and its interaction with T2DM on pulmonary tuberculosis in Chinese Uygur population. Oncotarget 10.18632/oncotarget.19274 (2017).

11. H. Wang et al., Identification of PTPN22, ST6GAL1 and JAZF1 as Psoriasis Risk Genes Demonstrates Shared Pathogenesis between Psoriasis and Diabetes. experimental dermatology 10.1111/exd.13393 (2017).

12. W. Wang et al., Association Between Protein Tyrosine Phosphatase Non-Receptor Type 22 (PTPN22) Polymorphisms and Risk of Ankylosing Spondylitis: A Meta-analysis. Medical science monitor: international medical journal of experimental and clinical research 23, 2619–2624 (2017).

13. K. Hasegawa et al., PEST domain-enriched tyrosine phosphatase (PEP) regulation of effector/memory T cells. Science 303, 685–689 (2004).

14. J. Zhang et al., The autoimmune disease-associated PTPN22 variant promotes calpain-mediated Lyp/Pep degradation associated with lymphocyte and dendritic cell hyperresponsiveness. Nature genetics 43, 902–907 (2011).

15. A. M. Wallis et al., TRAF3 enhances TCR signaling by regulating the inhibitors Csk and PTPN22. Sci Rep 7, 2081 (2017).

16. Y. Wang et al., The autoimmunity-associated gene PTPN22 potentiates toll-like receptor-driven, type 1 interferon-dependent immunity. Immunity 39, 111–122 (2013).

17. C. J. Maine, J. R. Teijaro, K. Marquardt, L. A. Sherman, PTPN22 contributes to exhaustion of T lymphocytes during chronic viral infection. Proceedings of the National Academy of Sciences of the United States of America 113, E7231–E7239 (2016).

18. T. Jofra et al., Extrinsic Protein Tyrosine Phosphatase Non-Receptor 22 Signals Contribute to CD8 T Cell Exhaustion and Promote Persistence of Chronic Lymphocytic Choriomeningitis Virus Infection. Frontiers in immunology 8, 811 (2017).

19. R. J. Brownlie et al., Resistance to TGFbeta suppression and improved anti-tumor responses in CD8(+) T cells lacking PTPN22. Nat Commun 8, 1343 (2017).

20. R. J. Brownlie, D. Wright, R. Zamoyska, R. J. Salmond, Deletion of PTPN22 improves effector and memory CD8+ T cell responses to tumors. JCI Insight 5 (2019).

21. T. Vang et al., Autoimmune-associated lymphoid tyrosine phosphatase is a gain-of-function variant. Nature genetics 37, 1317–1319 (2005).

22. A. Gregorieff, J.-F. Cloutier, A. Veillette, Sequence Requirements for Association of Protein-tyrosine Phosphatase PEP with the Src Homology 3 Domain of Inhibitory Tyrosine Protein Kinase p50. Journal of Biological Chemistry 273, 13217–13222 (1998).

23. J. F. Cloutier, A. Veillette, Association of inhibitory tyrosine protein kinase p50csk with protein tyrosine phosphatase PEP in T cells and other hemopoietic cells. The EMBO Journal 15, 4909–4918 (1996).

24. C. J. Maine, K. Marquardt, J. Cheung, L. A. Sherman, PTPN22 controls the germinal center by influencing the numbers and activity of T follicular helper cells. Journal of immunology 192, 1415–1424 (2014).

25. J. A. Knipper, D. Wright, A. P. Cope, B. Malissen, R. Zamoyska, PTPN22 Acts in a Cell Intrinsic Manner to Restrict the Proliferation and Differentiation of T Cells Following Antibody Lymphodepletion. Frontiers in immunology 11 (2020).

26. M. J. Smyth, G. P. Dunn, R. D. Schreiber, “Cancer Immunosurveillance and Immunoediting: The Roles of Immunity in Suppressing Tumor Development and Shaping Tumor Immunogenicity”. (Elsevier, 2006), 10.1016/s0065-2776(06)90001-7, pp. 1–50.

27. L. Haas, A. C. Obenauf, Allies or Enemies—The Multifaceted Role of Myeloid Cells in the Tumor Microenvironment. Frontiers in immunology 10 (2019).

28. J. P. Böttcher, C. Reis E Sousa, The Role of Type 1 Conventional Dendritic Cells in Cancer Immunity. Trends in Cancer 4, 784–792 (2018).

29. S. Yang, W. Wei, Q. Zhao, B7-H3, a checkpoint molecule, as a target for cancer immunotherapy. International Journal of Biological Sciences 16, 1767–1773 (2020).

30. T. E. Hickey et al., The androgen receptor is a tumor suppressor in estrogen receptor– positive breast cancer. Nature Medicine 27, 310–320 (2021).

31. K. Fujita, N. Nonomura, Role of Androgen Receptor in Prostate Cancer: A Review. The World Journal of Men’s Health 37, 288 (2019).

32. P. S. Ramos, A. M. Shedlock, C. D. Langefeld, Genetics of autoimmune diseases: insights from population genetics. Journal of Human Genetics 60, 657–664 (2015).

33. R. Cubas et al., Autoimmunity linked protein phosphatase PTPN22 as a target for cancer immunotherapy. Journal for ImmunoTherapy of Cancer 8, e001439 (2020).

34. A. M. Van Der Leun, D. S. Thommen, T. N. Schumacher, CD8+ T cell states in human cancer: insights from single-cell analysis. Nature Reviews Cancer 20, 218–232 (2020).

35. C. G. Clemente et al., Prognostic value of tumor infiltrating lymphocytes in the vertical growth phase of primary cutaneous melanoma. Cancer 77, 1303–1310 (1996).

36. M. F. Krummel, J. P. Allison, CD28 and CTLA-4 have opposing effects on the response of T cells to stimulation. Journal of Experimental Medicine 182, 459–465 (1995).

37. L. S. K. Walker, Treg and CTLA-4: Two intertwining pathways to immune tolerance. Journal of autoimmunity 45, 49–57 (2013).

38. N. Bottini, E. J. Peterson, Tyrosine Phosphatase PTPN22: Multifunctional Regulator of Immune Signaling, Development, and Disease. Annual Review of Immunology 32, 83–119 (2014).

39. R. Bos, L. A. Sherman, CD4+ T-Cell Help in the Tumor Milieu Is Required for Recruitment and Cytolytic Function of CD8+ T Lymphocytes. Cancer research 70, 8368–8377 (2010).

40. H.-J. Kim, H. Cantor, CD4 T-cell Subsets and Tumor Immunity: The Helpful and the Not-so-Helpful. Cancer immunology research 2, 91–98 (2014).

41. N. Bottini, E. J. Peterson, Tyrosine phosphatase PTPN22: multifunctional regulator of immune signaling, development, and disease. Annu Rev Immunol 32, 83–119 (2014).

42. R. J. Brownlie et al., Lack of the phosphatase PTPN22 increases adhesion of murine regulatory T cells to improve their immunosuppressive function. Science signaling 5, ra87 (2012).

43. M. Li et al., The common, autoimmunity-predisposing 620Arg > Trp variant of PTPN22 modulates macrophage function and morphology. Journal of autoimmunity 79, 74–83 (2017).

44. H. H. Chang et al., PTPN22 modulates macrophage polarization and susceptibility to dextran sulfate sodium-induced colitis. Journal of immunology 191, 2134–2143 (2013).

45. X. Huang et al., VISTA: an immune regulatory protein checking tumor and immune cells in cancer immunotherapy. J Hematol Oncol 13, 83 (2020).

46. H. Dong et al., Tumor-associated B7-H1 promotes T-cell apoptosis: A potential mechanism of immune evasion. Nature Medicine 8, 793–800 (2002).

47. S. Turajlic, M. Gore, J. Larkin, First report of overall survival for ipilimumab plus nivolumab from the phase III Checkmate 067 study in advanced melanoma. Annals of Oncology 29, 542–543 (2018).

48. S. Pilon-Thomas, A. Mackay, N. Vohra, J.J. Mulé, Blockade of Programmed Death Ligand 1 Enhances the Therapeutic Efficacy of Combination Immunotherapy against Melanoma. The Journal of Immunology 184, 3442–3449 (2010).

