## Supplemental Information for "Pro-autoimmune allele of tyrosine phosphatase, PTPN22, enhances tumor immunity"

**Supplemental Figure 1. Lymphocytes are needed to control tumors in both PEP-WT and PEP-619WW mice.** A) Breeding schematic used to create Rag1$^{-/-}$ mice with PEP-619WW expression. B16-OVA growth curve in B) Rag 1$^{-/-}$ PEP-WT individual mice, C) Rag 1$^{-/-}$ PEP-619WW individual mice. D) Average growth with SEM shown. N=5 for each genotype. T test with Welch's Correction at each time point was done to determine any significant differences between genotypes.

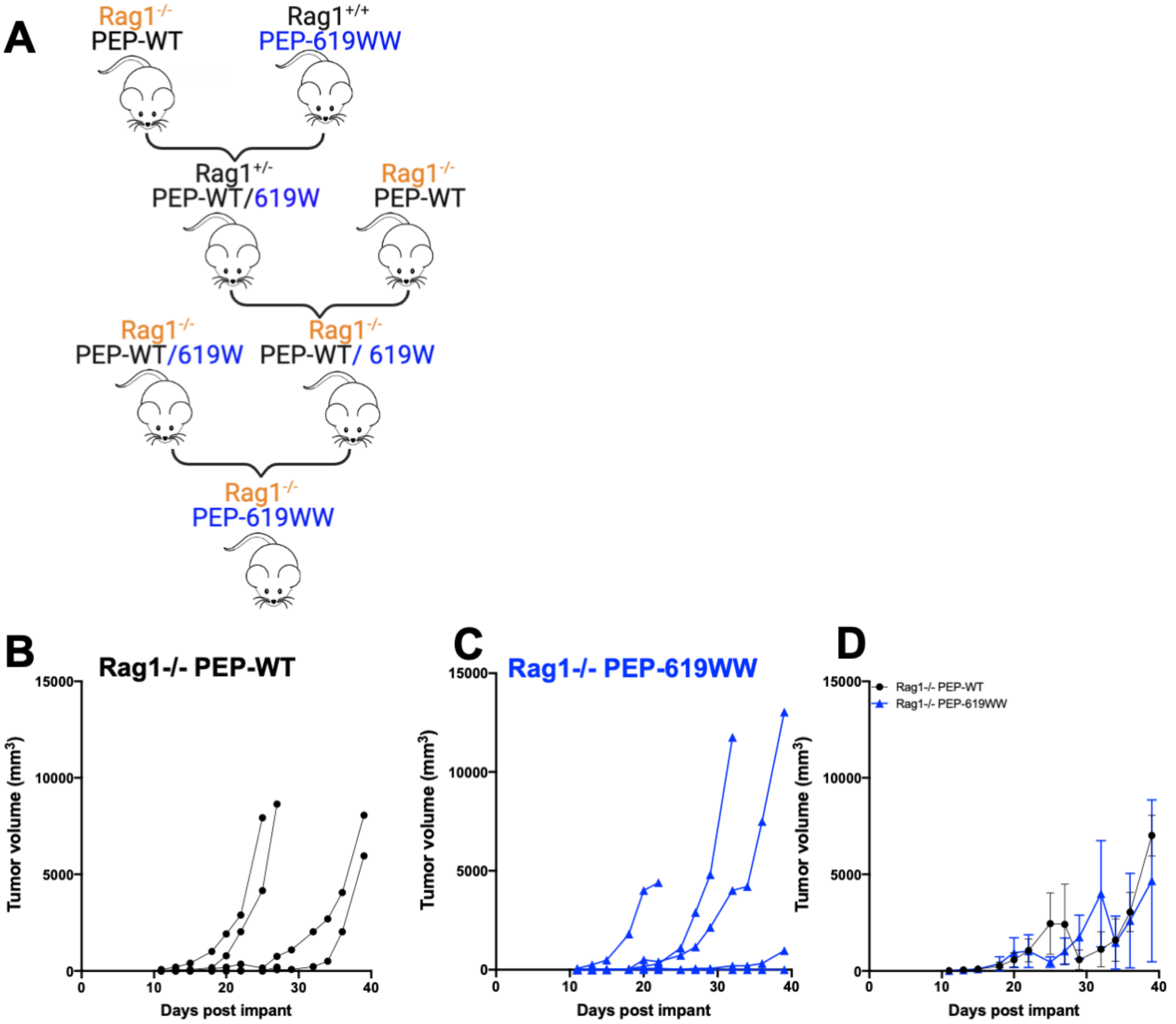

**Supplemental Figure 2. Nanostring GeoMX DSP Selected Regions of interest at CD45 dense regions.** A) Individual regions for each genotype (12 for PEP-WT and 12 for PEP-619WW mice). B) Example of 1/3 tumor sections for each genotype with regions of interest outlined in white. N=3 tumors per genotype.

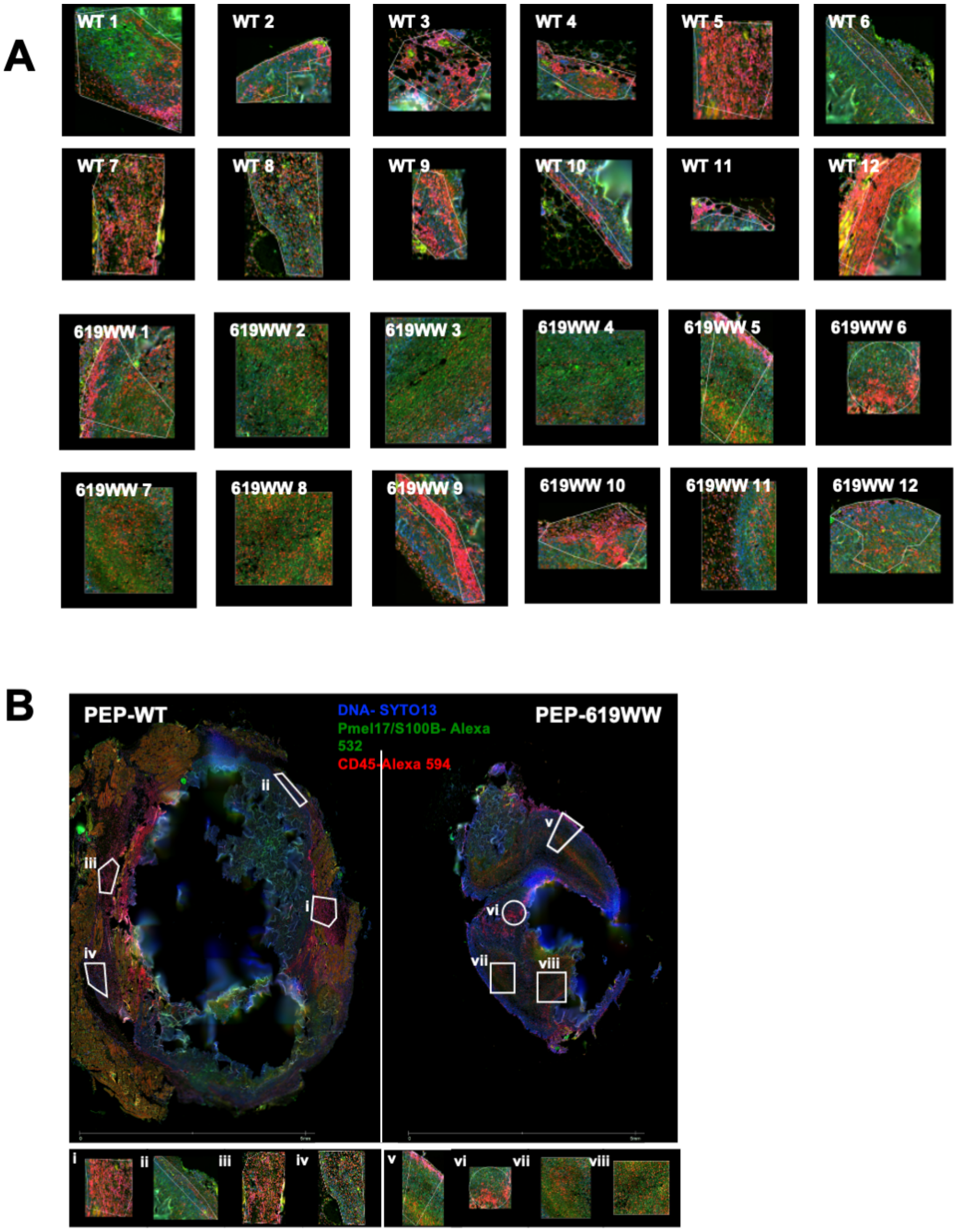

**Supplemental Figure 3. Non detectable difference in splenic Treg count between WT and PEP-619WW mice.** Spleens were harvested from naïve PEP-WT and PEP619WW mice. Absolute number Tregs in spleen with SEM shown. Gating strategy: Lymphocyte>Single cellx2> Live> CD45+> CD3+ CD19-> CD8a- CD4+>FoxP3+ CD25+. Significance determined via T Test with Welch's correction.

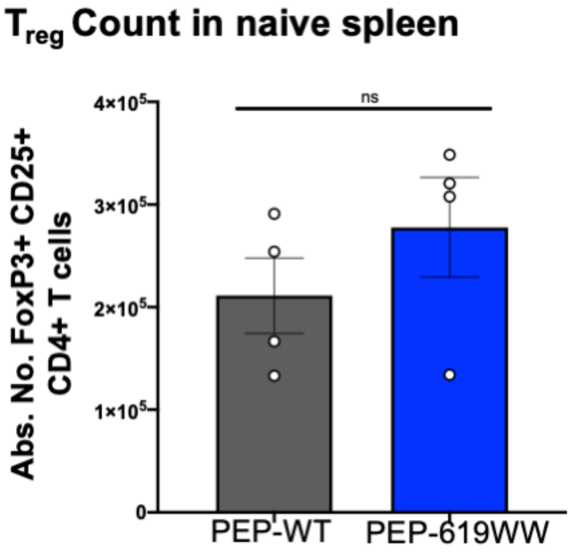

**Supplemental Figure 4. Genotyping PEP-619WW mice.** Amplified region of *Ptpn22,* which encompasses mutation sites responsible for the Arg→ Trp amino acid substitution at position 619, is digested with restriction enzyme BspEI. This site is only present in mice with the mutation. Digested fragments are run on an agarose gel and visualized. PEP-WT mice have a single band greater than 500 bp. PEP-619WW mice have 2 band less than 500bp each.

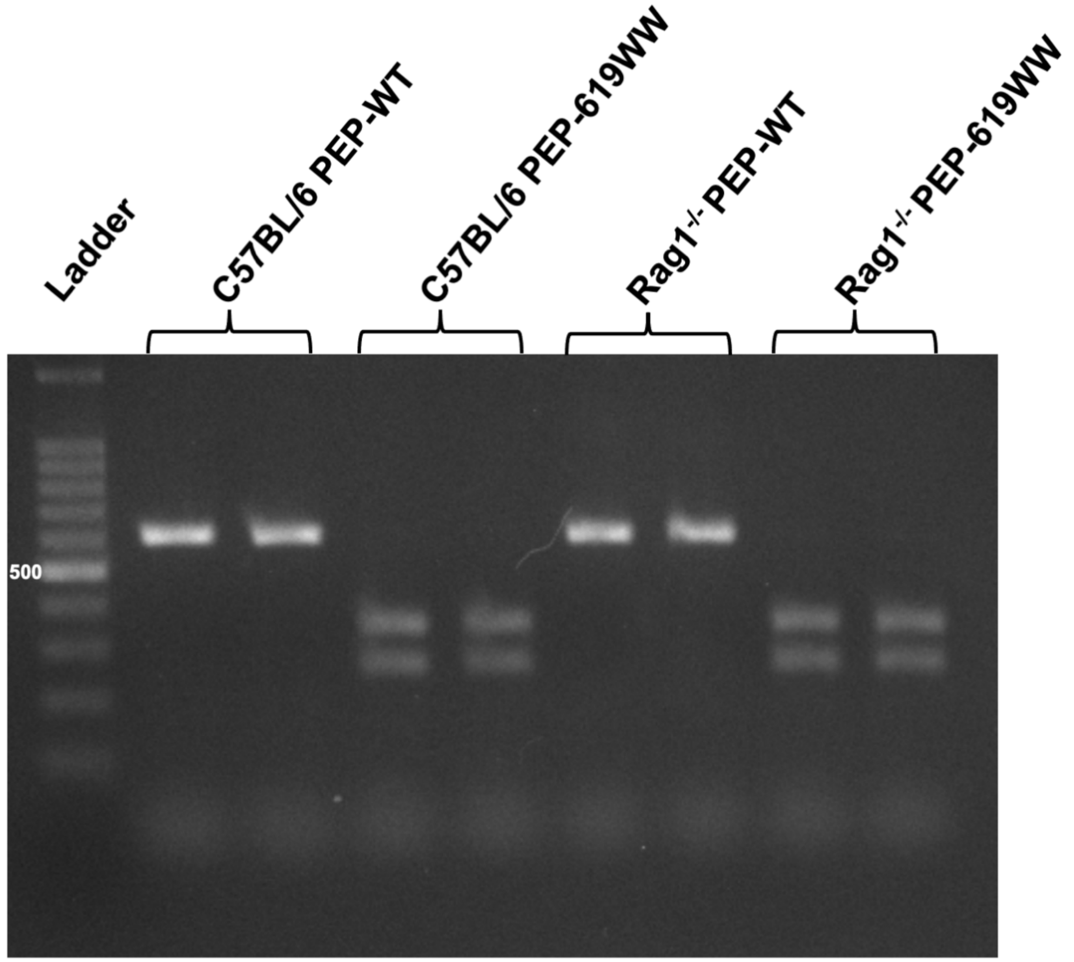
